# PeTFlow: Flow-Matching Based All-Atom Peptide Design through Side-Chain State Tokenization

**DOI:** 10.64898/2026.10.09.757799

**Authors:** Qirui Deng, Mingyuan Xu, Hongming Chen, Jinping Lei

## Abstract

Peptide offers a powerful modality for targeting protein-protein interfaces, yet their computational design remains challenging because sequence identity, backbone geometry and side-chain packing must be optimized simultaneously within a three-dimensional pocket constrained environment. Here we present PeTFlow, a pocket-conditioned all-atom peptide co-design framework that rationalizes this constrained design problem through discrete side-chain state tokenization. In PeTFlow, each residue is represented as a chemically compatible state token, enabling unified modeling of peptide sequence, conformation and all-atom interface geometry. PeTFlow achieved superior generation quality on peptide-design benchmarks and can further be applied to therapeutically relevant peptide drug design.

## Introduction

Peptide has emerged as an attractive therapeutic modality occupying a unique chemical space between small molecules and macromolecular biologics. Their intermediate size, conformational flexibility, and chemically diverse side chains enable them to recognize extended and relatively shallow protein surfaces that are often difficult to target with small molecules.^1^ These advantages have supported the broad clinical translation of polypeptides therapeutics, with more than 100 peptide drugs approved or marketed for the treatment of infection, diabetes, cancer, osteoporosis, multiple sclerosis, cardiovascular disease, chronic pain, and other disorders.^2,3^

Experimental peptide discovery has mainly relied on medicinal chemistry guided optimization and high-throughput library screening.^4,5^ Medicinal chemistry approaches commonly start from fragments of known binding proteins, endogenous ligands, or signaling peptides, followed by truncation, alanine scanning, residue substitution, and chemical modification to identify and optimize key binding determinants.^6-8^ However, these strategies depend strongly on existing binders and previously identified interaction motifs, limiting their ability to discover fundamentally new sequences and binding modes. Meanwhile, phage display and mRNA display enable the screening of large peptide libraries and have become powerful tools for identifying target-binding peptides.^9,10^ Nevertheless, these methods require substantial experiment effort and are sensitive to library composition, selection conditions, and subsequent hit validation and optimization.^11,12^ Hence, efficient computational methods are needed to explore peptide sequence and structural space more directly for accelerating the discovery of peptide binders with desired properties.

Among these methods, artificial intelligence has become an increasingly powerful framework for peptide design because of its ability to learn complex nonlinear relationships among peptide sequence, structure, and function from large biological datasets.^13-16^ The sequence-structure co-design methods, including DiffPepBuilder^17^ and PPFlow^18^, jointly generate peptide backbone geometries and amino-acid sequences. However, side-chain coordinates are not explicitly co-generated with the backbone and sequence in these methods. Recent all-atom co-design models primarily use either torsion-based or latent-space representations for side-chain conformations. Torsion-based methods such as PepFlow^19^ and ProBayes^20^ parameterize side-chain conformations using residue-dependent χ torsions, providing a compact representation but requiring amino-acid-dependent numbers of torsions and validity masks. Latent-space methods like PepGLAD^21^ and UniMoMo^22^ compress variable-size residues or molecular blocks into fixed-dimensional continuous representations, but the final all-atom structure depends on both latent-space sampling and decoder reconstruction. A unified representation that couples amino-acid identity with compatible side-chain conformations, while supporting explicit atom-level generation, remains lacking.

Here we introduce PeTFlow, a pocket-conditioned all-atom peptide co-design framework based on discrete side-chain state tokenization. Firstly, a VQ-VAE encodes each residue type and its side-chain conformation into a fixed-dimensional discrete token. Then, a residue-level generative model jointly samples peptide backbone frames and residue-state tokens under target-pocket conditioning. Finally, a VQ-conditioned atom-level generator produces complete side-chain coordinates while preserving the generated backbone. The performance of PeTFlow was evaluated on known peptide design benchmarks together with a series of baseline models, and the results demonstrated that PeTFlow achieves superior performance comparing with the chosen baselines.

## Results

### Overview of PeTFlow

PeTFlow is composed of three core modules: a VQ-VAE based side-chain state tokenizer, a Backbone–VQ Generator and a Side-chain generator (**Figure 1**). The VQ-VAE tokenizer compresses each residue type and its compatible side-chain conformation into a discrete residue-state token. This establishes a unified 1024-token vocabulary that represents residue-specific side-chain states, allowing PeTFlow to learn distributions over coupled residue–conformation states. The Backbone–VQ generator samples residue frames together with VQ tokens under pocket conditioning, and uses geometry-aware Invariant Point Attention Module (IPA) to jointly predict backbone frames and VQ token distributions, enabling generation of peptide backbone geometry and residue-state information for side-chain conformation. In the last step, the Side-chain generator firstly determines the representation of pocket-compatible backbone and side-chain state and then decode it into a complete all-atom peptide structure. By doing in this way, the generated backbone and VQ tokens from prior steps are converted into explicit all-atom coordinates in the protein pocket.

**Figure 1.**
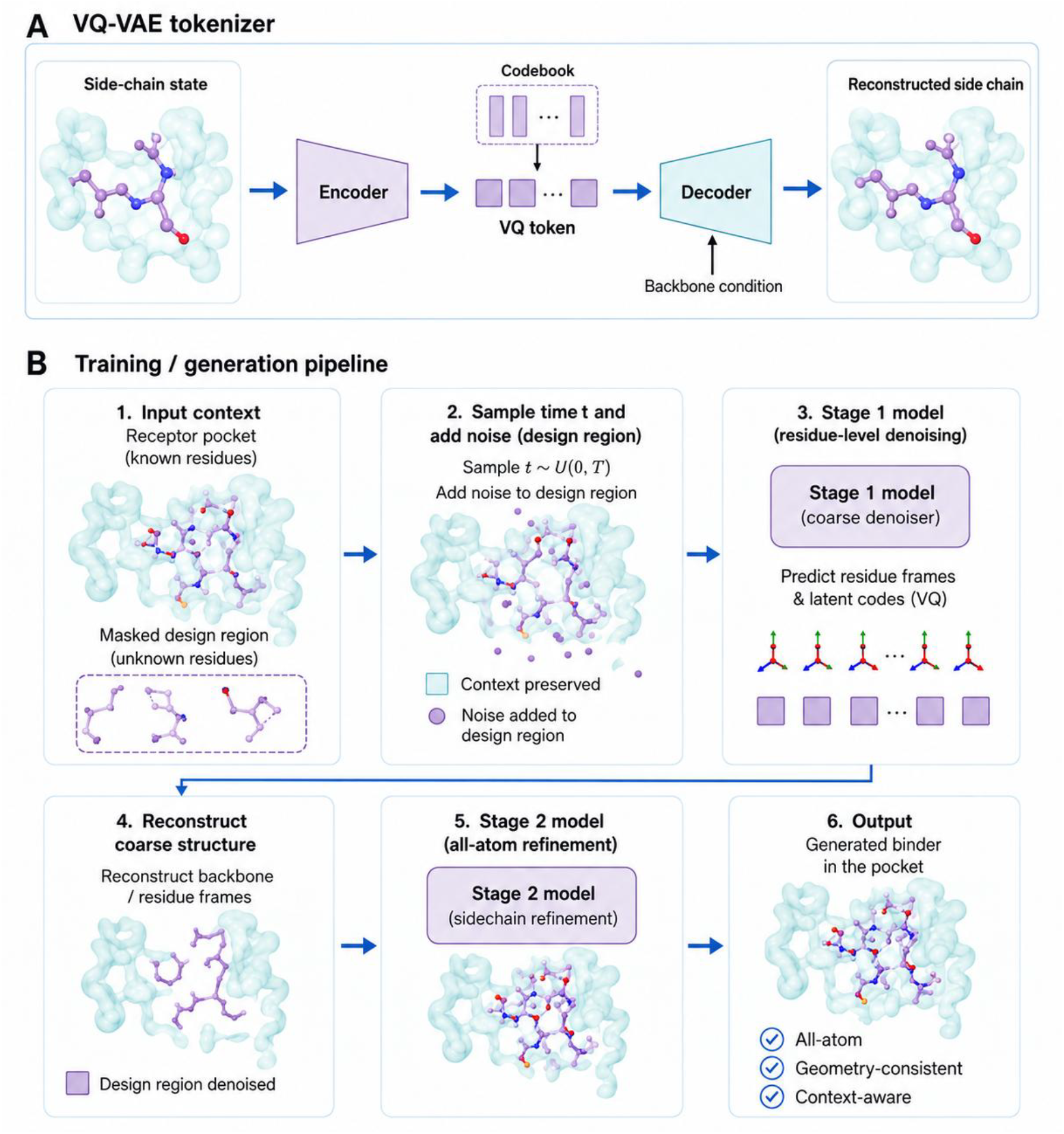
Overview of PeTFlow framework for pocket-conditioned all-atom peptide design.

To train the Backbone–VQ and Side-chain Generators of PepFloken, we first constructed a peptide-like protein fragment dataset containing 67,535 complex structures for pretraining of these two generators to learn diverse backbone structures, and then construced a peptide-protein complex dataset containing 10,184 structures for generators’ fine-tuning for peptide binder design. For side-chain state representation, the VQ-VAE tokenizer was trained on protein monomer structures from CATH 4.4.0 database^23,24^ to learn diverse side-chain conformations.

Due to the importance of VQ tokens in representing residue-level side-chain states for subsequent side-chain generation, we first evaluated whether the learned codebook could effectively capture and reconstruct side-chain conformation information. On the CATH monomer test set, the VQ-VAE achieved a codebook utilization of 99.5% (1019/1024 tokens) and a side-chain dihedral reconstruction MSE of 0.0925.

### PeTFlow achieves robust performance on target-specific peptide co-design

To evaluate the target-specific peptide co-design ability of PeTFlow, we compared it with two representative peptide-design models, PepFlow and PepGLAD, and two general protein-design frameworks, RFdiffusion and BoltzGen. For each of the 221 test complexes, all methods generated 40 same-length peptide candidates, which were evaluated in terms of backbone recovery, interface quality, sequence recovery, amino-acid composition and Rosetta interface energy.

Overall, PeTFlow achieved stronger native-like likeness than the peptide-generation baselines and remained competitive with the larger-scale protein-design models. As shown in **Table 1**, compared with PepFlow, PeTFlow reduced the overall Cα RMSD from 4.437 Å to 3.213 Å and improved DockQ from 0.379 to 0.424, PeTFlow also achieved higher amino acid recovery (AAR) than PepFlow and PepGLAD, indicating better recovery of peptide sequence and peptide–protein interface geometry.

**Table 1.** Evaluation on peptide co-design in native likeness. On each target, 40 candidates are generation for evaluation.

| Method | Native likeness |  |  |  |  |  |
| --- | --- | --- | --- | --- | --- | --- |
| | AAR (↑) | Cα RMSD (↓) | TM-score (↑) | DockQ (↑) | SSR(↑) | BSR(↑) |
| PeTFlow | **0.130** | **3.213** | <u>0.376</u> | **0.424** | **0.799** | **0.910** |
| PepFlow | <u>0.108</u> | <u>4.437</u> | 0.309 | <u>0.379</u> | <u>0.791</u> | 0.842 |
| PepGLAD | 0.067 | 4.922 | 0.274 | 0.299 | 0.755 | <u>0.899</u> |
| RFdiffusion | 0.074 | 5.129 | 0.353 | 0.170 | 0.663 | 0.558 |
| Boltzgen | 0.101 | 5.062 | **0.385** | 0.287 | 0.744 | 0.802 |

We next examined whether the performance of PeTFlow was affected by peptide length and secondary structure. PeTFlow showed clear advantages on short and medium peptides, achieving lower Cα RMSD, higher DockQ and TM-score than most baseline methods (**Figure 2A**). These improvements were statistically significant for short peptides and remained evident for medium-length peptides. For long peptides, however, the performance gap among peptide-generation methods became smaller. PeTFlow performed strongly on complexes with native loop peptides, and showed favorable performance on complexes with native strand-like peptides (**Figure 2B**). For complexes with native helical peptides, PeTFlow achieved backbone recovery comparable to the strongest baselines while maintaining improved interface quality. These results suggest that PeTFlow can recover native-like peptide binding modes across targets with different native peptide conformations.

**Figure 2.**
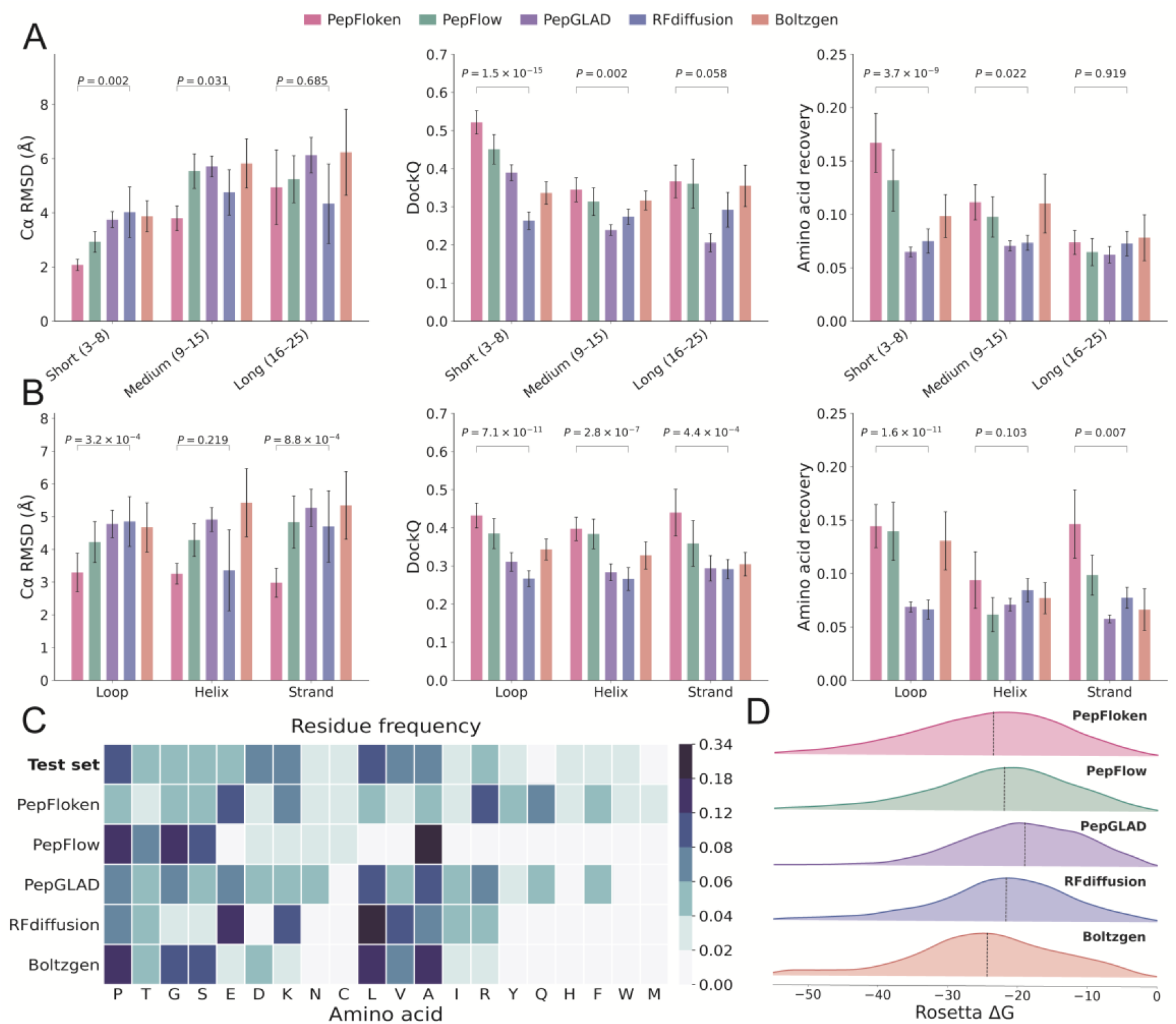
Performance of PeTFlow for target-specific peptide co-design. (A) Performance of PeTFlow and baseline methods on Cα RMSD, DockQ and AAR metrics across peptide-length groups. (B) Metrics of different models on the test set categorized by secondary structure including loop, helix and strand. (C) Residue frequency of the test set and peptides generated by different methods. (D) Distribution of Rosetta interface energy ΔG for generated peptide–protein complexes.

In addition to structural recovery, we evaluated the amino-acid composition of the generated peptides. The residue-frequency analysis showed that PeTFlow broadly preserved the amino-acid usage pattern of the test set and avoided severe collapse to a restricted subset of residue types (**Figure 2C**). This trend was particularly evident when compared with PepFlow which strongly enriched small and flexible residues such as Ala, Pro and Gly, and RFdiffusion that showed marked enrichment of Leu and Glu. In contrast, PeTFlow produced a more balanced amino-acid distribution and avoided the near-exclusion of specific residues observed in some baseline models, indicating PeTFlow can explore a broader peptide sequence space without collapsing to a narrow set of preferred amino acids.

Finally, we assessed the energetic plausibility of the generated peptide–protein complexes using Rosetta interface energy. PeTFlow generated favorable interaction-energy distributions, outperforming PepFlow, PepGLAD and RFdiffusion, although Boltzgen showed slightly lower Rosetta ΔG (**Figure 2D**). Together, these results demonstrate that PeTFlow robustly generates target-compatible peptide binders with improved backbone recovery, interface quality, balanced amino-acid composition and favorable interaction energy across diverse peptide classes.

## Conclusion

In this work, we introduced PeTFlow, a pocket-conditioned all-atom peptide co-design framework that integrates peptide backbone generation, amino-acid identity prediction and side-chain conformational modeling through discrete residue-state tokenization. By replacing explicit residue-dependent side-chain torsion prediction with a VQ-based side-chain-state representation, PeTFlow provides a unified and fixed representation for all-atom peptide generation. PeTFlow showed strong performance in target-specific peptide co-design, and outperformed classical peptide-generation methods PepFlow/PepGLAD and larger-scale general protein-design frameworks Boltzgen/AnewOmni. These results make PeTFlow potentially a useful tool for peptide design in drug discovery projects

## Methods

### Model architecture

PeTFlow comprises three core components: a vector-quantized variational autoencoder (VQ-VAE)^25^ that learns and stores the discrete side-chain-state codebook, a Backbone– VQ Generator that jointly generates peptide backbone conformations and VQ tokens, and a side-chain generator that reconstructs all-atom side-chain structures.

The VQ-VAE converts the heterogeneous combination of amino-acid identity and residue-dependent side-chain torsion into a unified discrete representation. Specifically, the encoder Eϕ embeds the residue type α_i_ and corresponding torsion angles χ_*i*_into a continuous latent vector through a multilayer perceptron, which is subsequently quantified to its nearest codebook entry:

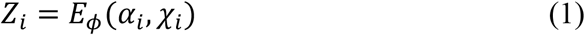

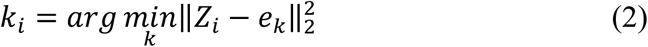

Conditioned on the quantified token and local backbone geometry, the attention-based decoder D_ξ_ reconstructs the residue type 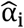 and side-chain torsion angles 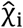:

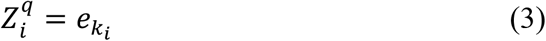

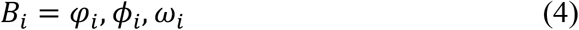

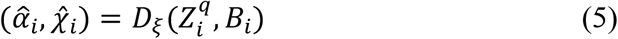

where *B*_*i*_ denotes the local backbone angles of phi (*φ*_*i*_), psi (*ϕ*_*i*_) and omega (*ω*_*i*_). The Backbone–VQ Generator jointly models peptide backbone frames and discrete side-chain-state tokens conditioned on the target protein environment. At time t, each peptide residue is represented by a translation vector Tt, a local rotation frame R_t_, and a discrete VQ-token state Z_t_. The generator predicts the corresponding clean backbone frames and VQ-token distribution:

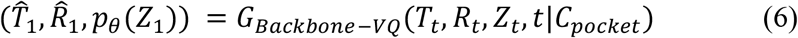

where C_pocket_ denotes the target protein environment, 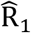 and 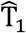 define the predicted residue-local backbone frames, and p_θ_(Z_1_) is the categorical distribution over the learned VQ codebook.

The side-chain generator converts the generated backbone–token representation into a complete all-atom peptide structure. Given the current side-chain atom coordinates state X_t_, the generated backbone frame of translation T and rotation R, VQ tokens Z, and time t, the generator predicts the corresponding clean side-chain atom coordinates:

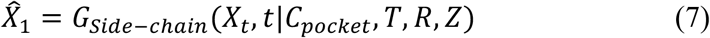

where 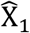 denotes the predicted ground-true coordinates state of the all-atom structure.

### Model Training

PeTFlow was trained in two major stages. First, the side-chain-state VQ-VAE was trained independently on monomeric protein structures to construct a discrete codebook that jointly represents amino-acid identity and side-chain conformational states. Then, the Backbone–VQ Generator and side-chain generator were optimized within the same training procedure.

The VQ-VAE is optimized using a joint reconstruction and vector-quantization objective:

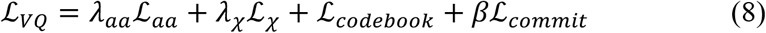

where *L*_aa_ is the cross-entropy loss for residue-type reconstruction, *L*_χ_ is a periodic torsion-angle reconstruction loss applied only to valid side-chain torsions, and the final two terms correspond to the codebook and commitment losses, respectively.

The second stage was optimized using a joint objective that supervises backbone generation, VQ-token prediction, sequence recovery, and all-atom side-chain generation. The backbone objective *L*_BB_ consists of the translation loss *L*_T_, rotation loss *L*_R_, and an auxiliary backbone geometry loss *L*_backbone_ . The translation and rotation terms supervise the predicted residue-frame origins and orientations, respectively, whereas the backbone geometry term constrains the reconstructed backbone atom positions and peptide-bond geometry:

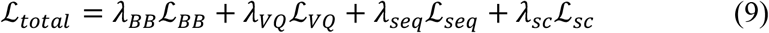

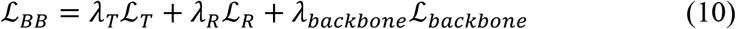

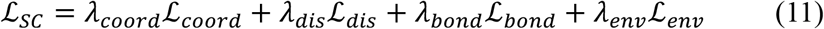

The VQ-token loss *L*_VQ_ supervises the categorical distribution over the learned residue-state codebook. In addition, the predicted VQ-token probabilities are projected to amino-acid probabilities and the resulting sequence distribution is supervised by *L*_seq_ . The side-chain objective *L*_sc_ combines coordinate- and geometry-level supervision. The coordinate loss *L*_coord_ directly penalizes deviations between predicted and native side-chain atom positions in residue-local coordinate frames. The intra residue distance loss *L*_dis_ preserves pairwise distances among side-chain atoms and between side-chain atoms and backbone anchors. The bond-length *L*_bond_ maintain chemically plausible local geometry. Finally, the environment-interaction loss *L*_env_ supervises distances between generated peptide side-chain atoms and nearby atoms in the surrounding protein environment, encouraging pocket-compatible side-chain placement. All side-chain losses are evaluated only for valid atoms in the generated peptide region.

## Code availability

The PeTFlow implementation will be released upon publication. The current prototype contains the residue-level flow model, VQ-token prediction module, VQ-conditioned side-chain decoder and atom-level attention refinement modules.

